# Evaporation and Focus Degradation Mitigation in In-Incubator Live Cell Imaging for Capacitance Lab-on-CMOS Microsystem Calibration

**DOI:** 10.1101/2025.06.14.659688

**Authors:** Y. Gilpin, C.-Y. Lin, M. Dandin

## Abstract

Lab-on-CMOS is an instrumentation technology that combines miniaturized bioanalytical hardware with complementary metal-oxide semiconductor (CMOS) electronics to provide integrated biosensing in a compact format. This paper focuses on a class of lab-on-CMOS systems that utilize capacitance sensing as a means to monitor cell cultures and track cell proliferation, as well as other cell life-cycle events. In this paradigm, changes in interfacial capacitance result from the activity of adherent cells at a bioelectronic interface. These changes are mapped to cell proliferation or life-cycle events using a ground-truth measurement such as live cell imaging from real-time microscopy. This paper identifies instrumentation challenges that arise from conducting these ground-truth measurements in a calibrated cell culture environment, *i*.*e*., when the lab-on-CMOS system is deployed inside a *CO*_2_ cell culture incubator. We show that autofocusing the microscopy column and provisioning the lab-on-CMOS with an immersion lid are two approaches that significantly improve the quality of live cell imaging ground-truth measurements over long periods.

## I. INTRODUCTION

The advent of lab-on-a-chip technology has given rise to new point-of-care capabilities, creating exciting opportunities for improving global health^1–3^. For example, in resource-limited settings, lab-on-a-chip technology opens the door to decentralized healthcare infrastructures where affordable and portable diagnostics tools provide rapid results without the need for specialized facilities or highly skilled technicians. As examples of applications that will benefit public health, lab-on-a-chip systems have been demonstrated in HIV diagnostics^4^ and in hematology^5,6^. Generally, they have been used in a wide variety of other assays with target analytes including cells, proteins, nucleic acids, or metabolites^7^. Lab-on-a-chip systems leverage small sample volumes and rapid processing to give high-fidelity results at a fraction of the typical cost. In sum, these capabilities illustrate the technology’s enormous potential for revolutionizing healthcare.

Another area in which lab-on-a-chip technology is expected to make advances is in enhancing current laboratory testing capabilities. In this approach, rather than emphasizing portability, the goal is to leverage inherent parallelization, rapid processing, and low sample volume features of lab-on-a-chip devices to enhance assay throughput and cost, and to provide novel capabilities in a traditional laboratory setting. For example, Chitale *et al*. showed a 96 well plate for in-incubator use and in which each well was provisioned with complementary metal-oxide semiconductor (CMOS) integrated circuits configured to perform impedance-based measurements^8^. Using more than a dozen plates, they screened 904 compounds against various cell types, demonstrating a new instrumentation technology for high-throughput action profiling and phenotypic drug discovery. Another notable example is from Hu *et al*.^9^; they showed a CMOS-based lab-on-a-chip platform for imaging and monitoring cells utilizing a range of sensing modalities. Once scaled, these chips will enable “wide-field monitoring of cells, colonies, or even whole tissues at a fraction of the size and cost of traditional microscopes^9^.”

Both of these examples fall under the lab-on-CMOS paradigm. In these devices, a CMOS chip is integrated in a microfluidic lab-on-a-chip, or generally, a CMOS chip is an integral part of the lab-on-a-chip. This approach is beneficial in locally integrating high spatial density sensing, signal processing, and control, providing a bioelectronic interface to monitor various aspects of a bio-specimen in real time. A key requirement in developing these lab-on-CMOS platforms includes calibrating the CMOS sensors using ground-truth data. For example, for a CMOS voltage-to-frequency pH sensor, calibration required benchmarking the sensor’s output against known pH values obtained with a reference instrument^10^.

In our own work, we have used imaging with upright bright-field optical microscopy to calibrate and train capacitance-sensing lab-on-CMOS platforms designed for in-incubator cell culture characterization^11,12^. When used for calibration, imaging correlates the measured capacitance with cell culture growth, particularly with cell numbers and cell number temporal evolution^11^. Once calibrated, the sensors can operate without imaging to provide an instantaneous measure of the number of cells in a region of interest (ROI) or a measure of the temporal evolution of the number of cells in the ROI. When used for training, imaging provides ground-truth data for a back-end algorithmic system that can recognize specific micro-trends in the measured time series capacitance data. As we have shown, with training, the algorithmic back-end can recognize the micro-trends directly from the time series capacitance data (*i*.*e*., with no imaging) and classify these micro-trends as mitosis or migration events^12^.

Figure 1 presents an overview of the capacitance-sensing lab-on-CMOS platform considered in this paper. Specifically, herein, we address the significant challenges encountered with in-incubator, upright, bright-field optical microscopy imaging when it is used to calibrate the capacitance-sensing lab-on-CMOS platform. We show that cell media evaporation dynamics are a major impediment to acquiring high-quality ground-truth images necessary for accurate cell counting via a computer-vision algorithm. We show two strategies that may mitigate this issue: (1) the use of a novel immersion lid with side ports for gas exchange and (2) the use of autofocusing. We provide a quantitative study assessing the performance of both methods, when they are used either alone or in combination. Section II describes the experimental methods used in this study. In Section III, we cover experimental results relating to capacitance sensing dynamics and image focus degradation and restoration via autofocusing, all in view of cell media evaporation. Section IV summarizes our findings and discusses the limitations of our study.

**FIG. 1.**
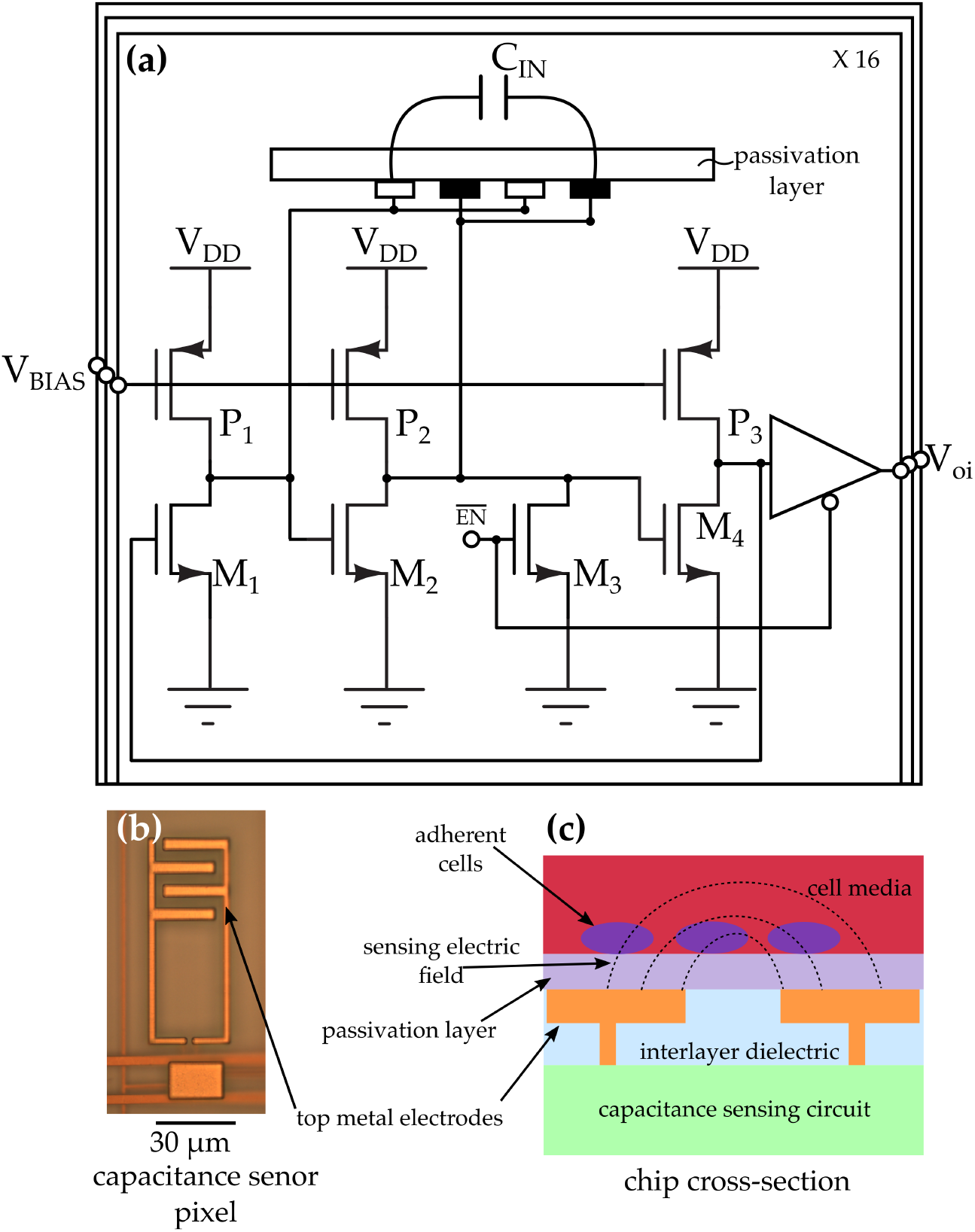
(a) Circuit schematic view of the CMOS capacitance sensor used in this study. The chip is a 4 × 4 capacitance sensor array where each pixel includes a three-stage oscillator circuit whose intermediate stage is connected to a passivated interdigitated electrode pair implemented in the CMOS process’ top metal layer. The oscillator’s output is buffered, and a pull-down switch (M3) is included to disable the pixel when it is not in use. An analog voltage (*V*_*BIAS*_) is used to set the base frequency of the oscillator. During operation, the output voltage of a pixel *i* is a digital signal *V*_*oi*_ whose frequency depends on the capacitance *C*_*IN*_ at the interface^13^. (b) Photomicrograph of one of the 16 electrodes of the test chip. (c) Cross-sectional sketch of the CMOS sensor chip around one electrode pair. We note that in this sketch, the electric field lines are not drawn to scale or with physical accuracy.

## II. EXPERIMENTAL METHODS

### A. Lab-on-CMOS Design and Packaging

Cell membranes typically exhibit surface charge density, which may be either positive or negative depending on the specific cell type^14,15^. This surface charge can create a cloud of counterions in the surrounding cell media^14^. Therefore, under electric field exposure, these counterions can shift tangentially along the cell membrane, generating dipole moments, which give rise to polarization effects^14^. The capacitive characteristics of cells thus arise in part because of these polarization effects^14^. In our CMOS capacitance sensors shown in Figure 1, the base frequency of the sensing oscillator is ~10 MHz. Therefore, when the oscillator is loaded, *i*.*e*., when polarization effects occur at the interface, the frequency slows. Because the oscillator’s frequency is inversely proportional to capacitance, there is a resulting increase in measured capacitance when loading occurs.

The sensing electrodes are designed as an inter-digitated pair of metal electrodes^16^. The electrodes are insulated using the CMOS chip’s passivation layer, and during operation, a portion of the resulting electric field between the two electrodes passes through the passivation layer and into the cell media above it. This allows the chip to detect cells at the interface. Generally, any perturbation at the interface may be interpreted by a circuit connected to the electrodes to provide insight into the activity at the interface. This makes integrated capacitance sensing a common technique for lab-on-CMOS devices used in live cell culture monitoring applications. Many circuit architectures have been demonstrated in integrated capacitance sensing^17^. The one featured in this work was a capacitance-to-frequency converter designed in a 0.35 *µm* CMOS process (see Figure 1). It uses a three-stage ring oscillator to map the measured capacitance at the interface to the frequency of a digital signal; the frequency is estimated by counting the number of edges registered by a digital counter during a fixed measurement period^11,13,18–21^.

One of the main challenges with integrated capacitance sensing is maintaining the electrical integrity of the chip while providing a means for an aqueous sample to be interfaced with the chip’s sensing electrodes. Several packaging approaches have been shown for achieving these two features. They include low-temperature co-fired ceramics, the use photo-polymers to encapsulate sensitive parts of the chip, photolitographically defined molds, or micromachined chip carriers^23–26^. Our approach (Figure 2) consisted of a die-on-PCB assembly where the CMOS die is attached and wirebonded to the PCB. The wirebonds are encapsulated with an impermeable epoxy (Duarpot 865, Cotronics Corp. Brooklyn, NY). The encapsulation process uses a dam to prevent the epoxy from reaching the sensor areas, leaving them exposed. After applying the epoxy, it is left to cure at room temperature for 24 hours. The assembly is then baked for 1 hour at 121^°^ C and subsequently at 177^°^ C for 5 hours^11,27^. This hard-baking step turns the epoxy from gray to an amber color. Following the die-on-PCB process, the assembly is fitted with a culture well and with a removable immersion lid.

**FIG. 2.**
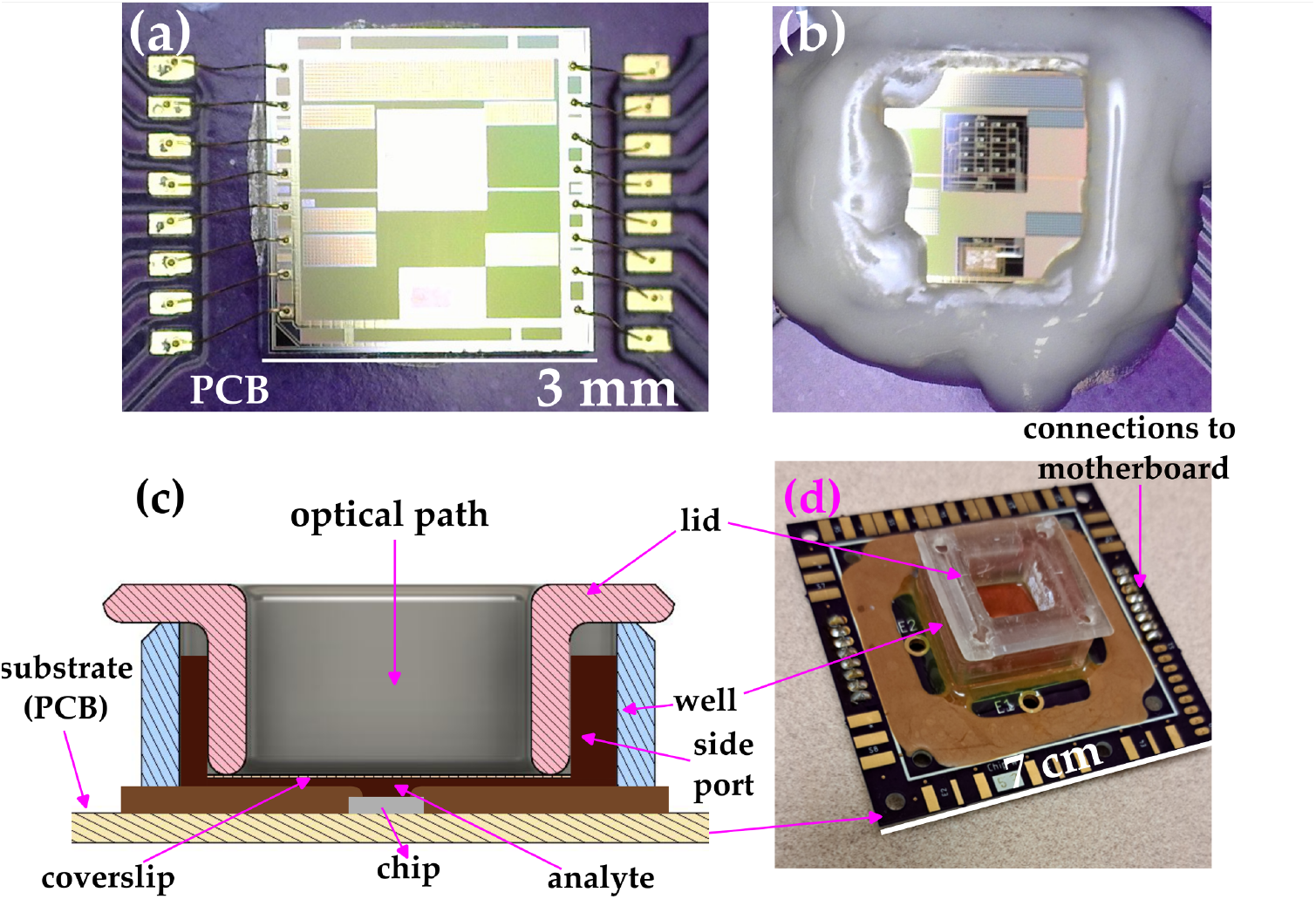
(a) Photomicrograph of the CMOS capacitance sensor mounted and wirebonded on a PCB. (b) Wirebond encapsulation with a biocompatible and impermeable epoxy. (c) Cross-sectional view of the immersion lid used in this study. (d) Photograph of the fully integrated lab-on-CMOS platform.

Figure 3 shows a diagram of the immersion lid designed for our experiments along with several other approaches used in the state-of-the-art. Our approach has several advantages over simple lids and over the dual well approach reported by Senevirathna *et al*^22^. Our lid reduces the volume of media in the optical path, which in turn reduces optical losses and sensitivity due to clouding. Furthermore, it is more resistant to evaporation because the optical path length is independent of small changes in media volume. Further, bubbles tend to float to the space on the side of the lid, away from the optical path and via the lid’s side ports. The immersion lid and the well were 3D printed using a Phrozen Sonic Mighty 8K 3D printer in NextDent Ortho Clear resin and washed using 100% ethanol for 5 minutes in an ultrasonic bath. Next, they were UV exposed for 10 minutes for further curing. At the 5 minute mark, the parts were rotated to improve the uniformity of the exposure. After UV curing, the parts were washed again in 100% ethanol for 5 minutes in an ultrasonic bath. A coverslip was then glued to the window opening using medical grade silicone (World Precision Instruments, Sarasota, FL). The well was also glued to the daughter board using the same silicone.

**FIG. 3.**
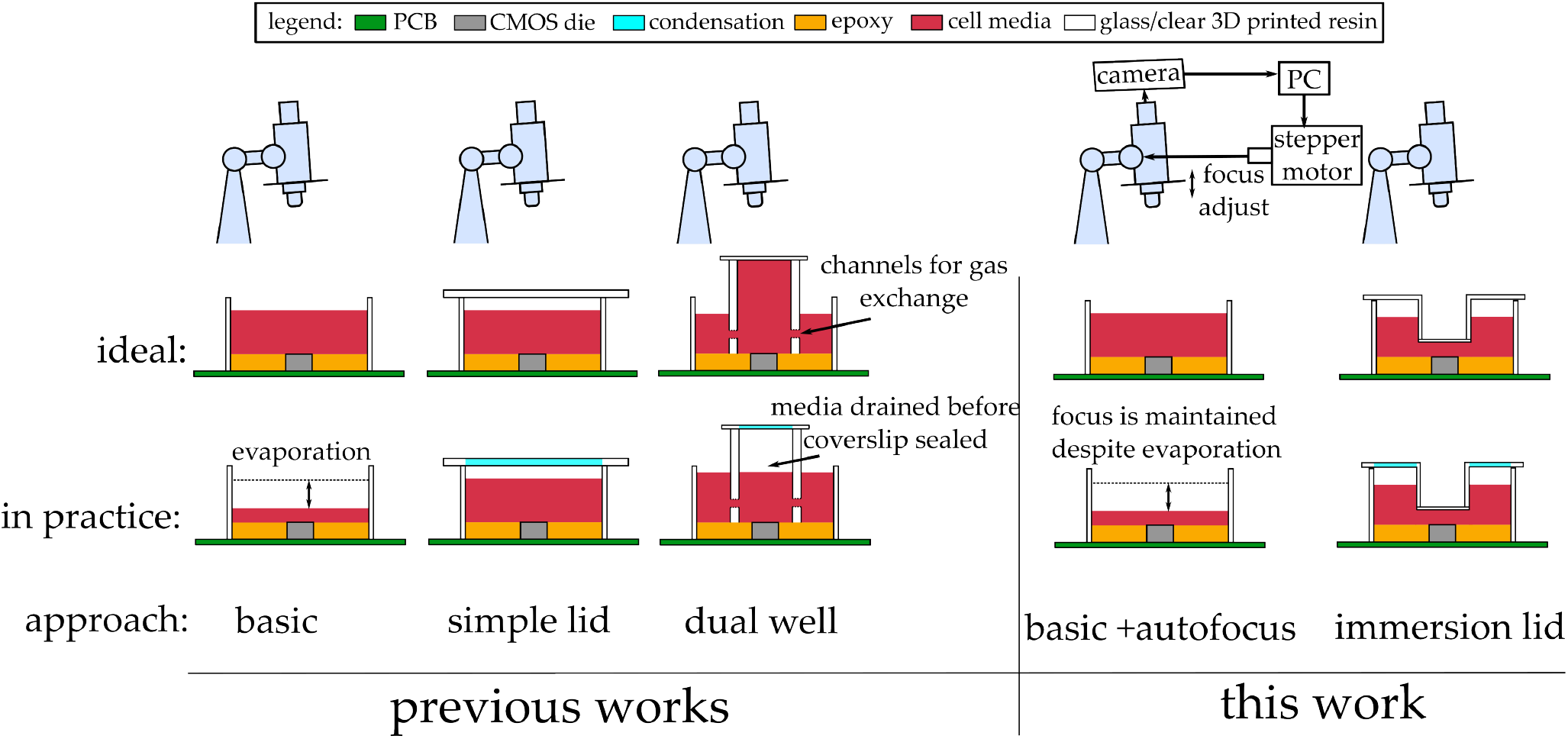
Comparison of upright microscope imaging techniques for live cell lab-on-CMOS cell culture. The basic method leads to blurry images due to evaporation. The simple lid also has blurry images due to condensation in the optical path. The dual well design featured in ref.^22^ mitigates the condensation issue but it can be difficult to use because the cell media often flows from the inner well to the outer well before the cover slip can seal. This work features two techniques; in the first, we use a custom autofocusing procedure combined with the basic lab-on-CMOS imaging setup, and in the second, we use a custom immersion lid.

### B. Experimental Setup

Our experimental setup included a Thermo Scientific Forma Series 3 water-jacketed cell culture incubator set and maintained at 37^°^ C and 5% *CO*_2_ (Figure 4). We used an in-chamber microscope to image the cell cultures under study. The microscope was a MicroZoom II bright-field up-right microscope from Bausch and Lomb, with an 5X objective and an approximate working distance of 2 cm. The microscope was equipped with an 18 mega-pixel C-mount digital camera (MU1803, Amscope, United Scope, CA). Images acquired from the camera were sent to a PC outside the incubator using a flat USB cable. This cable allowed us to maintain the incubator’s seal for the duration of our experiments. A humidity pan was placed inside the incubator to maintain a relative humidity close to 100% in the chamber. We note that without the humidity pan, the evaporation rates observed would be much higher than the ones reported below.

**FIG. 4.**
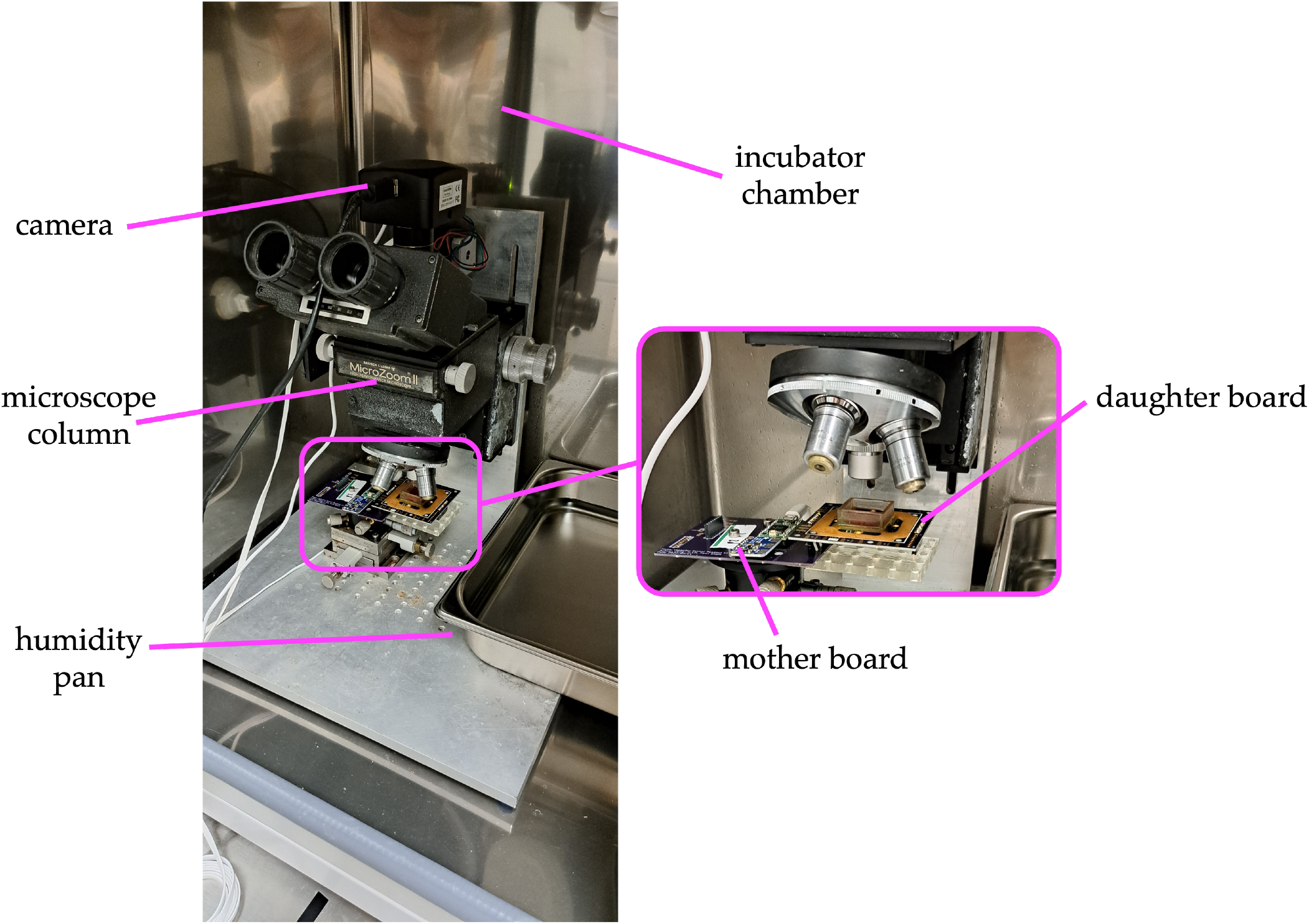
Photograph of the experimental setup configured for simultaneous live cell imaging and real-time lab-on-CMOS capacitance measurements.

The interface between the CMOS die and the PC was implemented with a daughter board mounted on a mother board. The latter consisted of a Teensy 3.2 (PJRC Sherwood, OR) microcontroller (32b, 72MHz) which generates control signals for the chip, the 3.3V power supply, and read the output buffer over an I^2^C. The clock needed to operate the chip was generated using an Adafruit (Brooklyn, NY) Si5351A clock generator module programmed at 4 MHz. The mother board, with the daughter board mounted thereon, was placed on a positioning stage under the objective.

### C. Cell Culture, Device Cleaning, and Sterilization

All the experiments in our study used RAW 264.7 murine macrophages (ATCC Manassas, VA). They were cultured in a cell culture flask (ThermoFisher, Waltham, MA) and in DMEM (ATCC) with 10% FBS (ATCC) and 1% PSG (Corning Inc. Corning, NY) media. To subculture the cells, the cell media was first aspirated and replaced by 5 mL of new media. The cells were lifted from the bottom of the culture flask using a cell scraper (VWR Radnor, PA). A 2:3 dilution was then used to subculture the cells in a new flask, at which point the cells were then plated on the CMOS chip at a density of 300,000 cells/mL in a total volume of 1.75 mL. All experiments were run for at least 36 hours but no longer than 40 hours.

After each experiment, the media in the cell culture well was discarded, and the well was sprayed with 70% ethanol. Subsequently, Tergazyme (Alconox, White Plains, NY) was used to clean any dead cells or other organic residue left on the CMOS die’s surface. The device was placed in a sonicated water bath for 5 minutes to further enhance the effects of the Tergazyme. After sonication, the Tergazyme was aspirated, and the well was sterilized by spraying it with 70% ethanol. Lastly, the device was washed 3 times with sterile PBS (ThermoFisher). As for the lid, spraying it with 70% ethanol and washing it 3 times with PBS was sufficient; cleaning it did not require a Tergazyme and sonication cycle because cells do not adhere on the lid’s inverted surface. Following this cleaning and sterilization process, the chip was ready for use in another experiment.

## III. RESULTS

We now report on capacitance measurement results, and we present our characterization of the autofocusing algorithm’s performance. To study capacitance dynamics, we measured the temporal evolution of the change in capacitance with cell media, with and without an immersion lid, with and without cells, and simultaneously with either fixed or automatic focusing on the microscope. For image quality assessment, the microscope was manually focused at the start of each experiment. This position was recorded and served as a reference position. At any given time during an experiment, the microscope column could be actuated back to the reference position in order to acquire a *fixed focus* image. In contrast, we acquired *autofocused* images by first computing a focus measure with the autofocusing algorithm and subsequently actuating the microscope column to a new position that maintained the computed focus measure. We benchmarked our results against a computer vision-based cell counting algorithm. Taken together, our results highlight the interplay between media evaporation, the effects of the lid, and focusing, and they further show how the sensor readings are influenced by these factors.

### A. Capacitance Dynamics

We measured the capacitance dynamics with media only on the chip, and we measured them with cells present in the media; for each scenario, we studied the dynamics with and without an immersion lid. Figure 5 shows the results of the media-only experiments. We pippetted 1.75 mL of cell media on the chip and placed it in a cell culture incubator for at least 36 hours. The experiments were repeated four times, and the measured capacitance data were averaged and subsequently fit. We found that capacitance decreased linearly with time, indicating a constant rate of media evaporation, whether a lid was used or not.

**FIG. 5.**
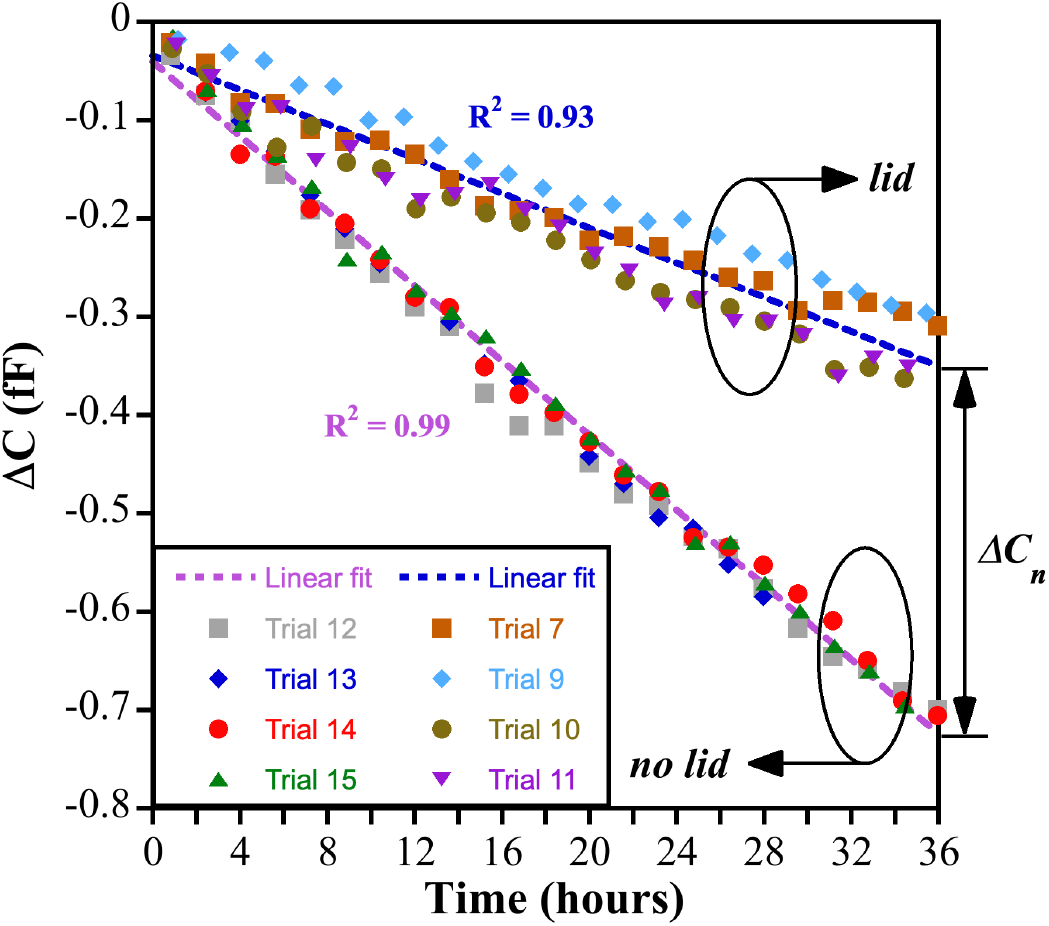
Media-only experiments. Eight trials were run, each lasting at least 36 hours. The data were sampled every 1.5 second but decimated for ease viewing. In four of the trials, we used an immersion lid, and in the other four, no lid was used. Here, each dataset (*i*.*e*., each trial) is the average of the measured capacitance change at all of the 16 sensors on the chip. For each condition (lid/no lid), the data were averaged across the four trials, and a linear fit was used to describe the resulting average (lid: *R*^2^ = 0.93, no lid: *R*^2^ = 0.99).

However, as expected, the capacitance signal decayed slower when a lid was used. At the 36-hour mark, the factor Δ*C*_*n*_, which denotes the difference in the capacitance change between the lid case and the the no lid case, was ~ −400 aF. This metric offers a quantitative assessment of the lid’s effect on media evaporation. More importantly, however, the results of Figure 5 show that while the lid slows down evaporation, it does not completely eliminate it. This is in fact a desirable result because it means that the lid does allow gas exchange between the media and the ambient environment; if that were precluded, cells would not be able to thrive in the media.

Turning now to Figure 6, we show the measured capacitance dynamics from experiments where cells were added to the media. Six experiments were run in total, each for at least 36 hours. Of the six experiments, three were run with an immersion lid, and three were run without one. Capacitance measurements started after an initial period to let the cells sediment on the sensor’s surface. The resulting capacitance dynamics were markedly different from those shown in Figure 5. For example, without a lid, the measured capacitance showed a noticeable decrease from 0 to 16 hours and a slightly increasing trend thereafter (see Figure 6(a)). With the lid, the response was more muted in the beginning, but the data also showed a slight increase in the second half of the measurement period (see Figure 6(b)).

**FIG. 6.**
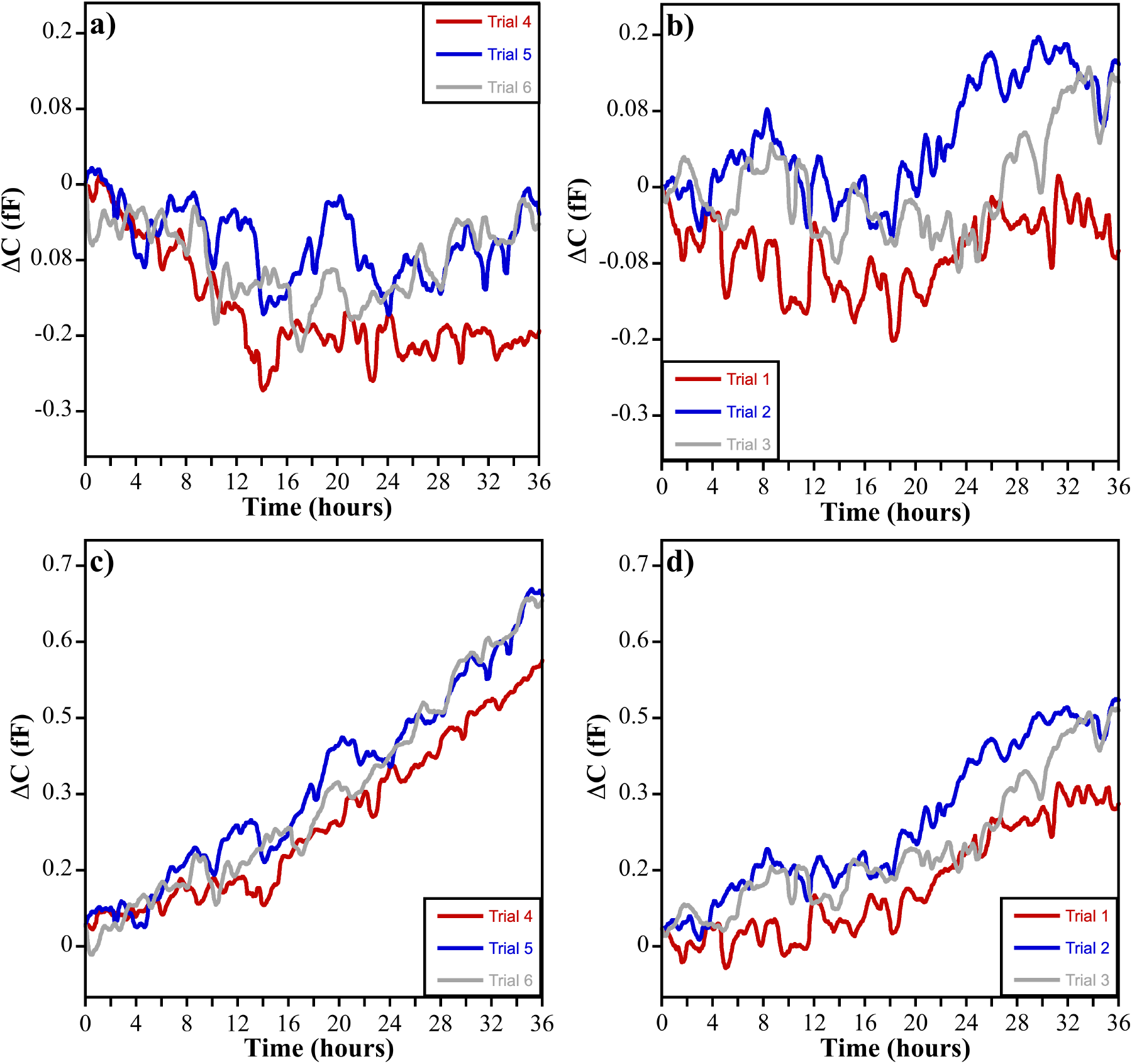
(a, b) Cells and media experiments. (a) Three trials were run without a lid, with 1.75 mL of cell media and cells were plated at a density of 300,000 cells/mL. (b) Another three trials were run with an immersion lid on the chip, with 1.75 mL of cell media and cells were plated at a density of 300,000 cells/mL. (c, d) Evaporation-compensated capacitance data for three trials conducted each without a lid (c) and for another three conducted with a lid on the chip (d).

Considering the no-lid case, the beginning downward transient indicates that evaporation dominates, as observed in the media-only case. However, as the cells divide and increase in number, the downtrend caused by media evaporation is countered. The muted initial response in the lid case means that the lid slows the evaporation rate; this is consistent with the results of Figure 5. Therefore, for both cases, we argue that the overall capacitance response is a composite effect that includes a media evaporation component and a cell division-induced component. Furthermore, the results of Figure 5 imply that the rate of change of capacitance is constant when considering the evaporation of the media. For example, for the no lid case, the rate was −20 aF/hour, and for the lid case, the rate was −8 aF/hour. This means that capacitance data related to cell culture growth can be compensated by subtracting the evaporation rate from the capacitance data shown in Figure 6. The results of this compensation are shown in Figures 6(c) and (d).

### B. Focus Dynamics

Having described the capacitance dynamics and demonstrated a technique for compensating evaporation effects, we now turn to describing ground-truth image quality and describe the issues that arise in image acquisition. For quantitative comparisons, we used a focus measure to assess image quality; the higher this focus measure, the higher the image quality. Furthermore, the focus measure was used as a control parameter for the autofocusing algorithm.

We used a second derivative-based focus estimation method to calculate the focus measure^28^. While there are multiple methods for estimating focus^29–33^, we chose a second derivative-based approach because second derivative operators are well suited for passing high spatial frequencies typically associated with sharp edges. These characteristics (sharp edges) are prominent in the images collected in our application. Specifically, cell membranes appear as sharp edges, and the chip’s own features include rectangular metal routing lines that form dense patterns presenting a large number of sharp edges. Interestingly, as an aside, though the chip’s features are helpful for assessing focus, they present quite a challenge for the computer vision algorithm used for cell counting, as we shall discuss below.

To compute the focus measure, we used the Laplacian as our second derivative operator. The Laplacian was approximated using the mask *L* as shown in Equation 1. Further, values Γ(*m, n*) were computed using a convolution of the input image intensity *I*(*m, n*) with the mask *L*. The indices *m* and *n* ranged from 1 to *M* and from 1 to *N*, respectively, for an image of *M* × *N* pixels. The focus measure *F* was calculated using Equation 2, which is the statistical variance of the convolution of the image *I* with the Laplacian mask *L*. In Equation 2, the average 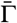 of the Γ(*m, n*) values was calculated using Equation 3.

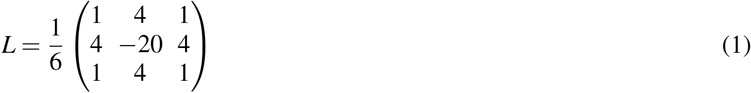

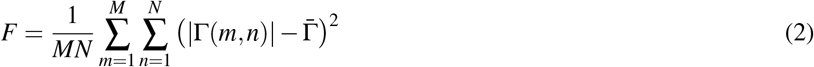

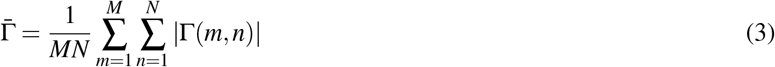

We studied focus dynamics under different conditions, *i*.*e*., we studied how the focus measure *F* varied with time in experiments where cells were loaded in the media, with or without a lid. The same volume and cell density used in the previous experiments were also used. Furthermore, images were taken at 5 minute intervals, and the focus measure was computed for each image and plotted as a function of time. At the beginning of each experiment, *i*.*e*., at t = 0, the microscope was focused manually, and the position was recorded by the control system. This position served as a reference. Image acquisition included taking an image at the reference position and taking another image at an auto-focused position. The latter was taken after actuating the microscope column to an auto-focused position according to the control algorithm. Therefore, for the same experiment, images in both focus settings could be acquired in order to study focus degradation over time but also to study the performance of the autofocusing algorithm.

Figure 7(a) shows measured focus dynamics for two experiments, one with a lid and one without a lid, when the microscope column is at the reference position. In this setting, the image is focused at the beginning of the experiment, and subsequent images for which the focus measure is computed are acquired with the microscope at that same position. The results show that when no lid is present, *F* degrades linearly. In contrast, with a lid, the focus degrades initially, but it levels off. Generally, the result shows that *F* is larger at all times when a lid is used, suggesting higher-quality images. Figure 7(b) shows experimental results when autofocusing is used. Without a lid, *F* remains constant. This suggests that the autofocusing algorithm can maintain the same image quality as that of the image acquired at the reference position. However, when a lid is used, focus degrades initially and subsequently levels off, similar to the experiment where no autofocusing was used. However, in contrast with the results of Figure 7(a), autofocusing does not provide a greater focus measure all throughout the experiment when the lid is used.

**FIG. 7.**
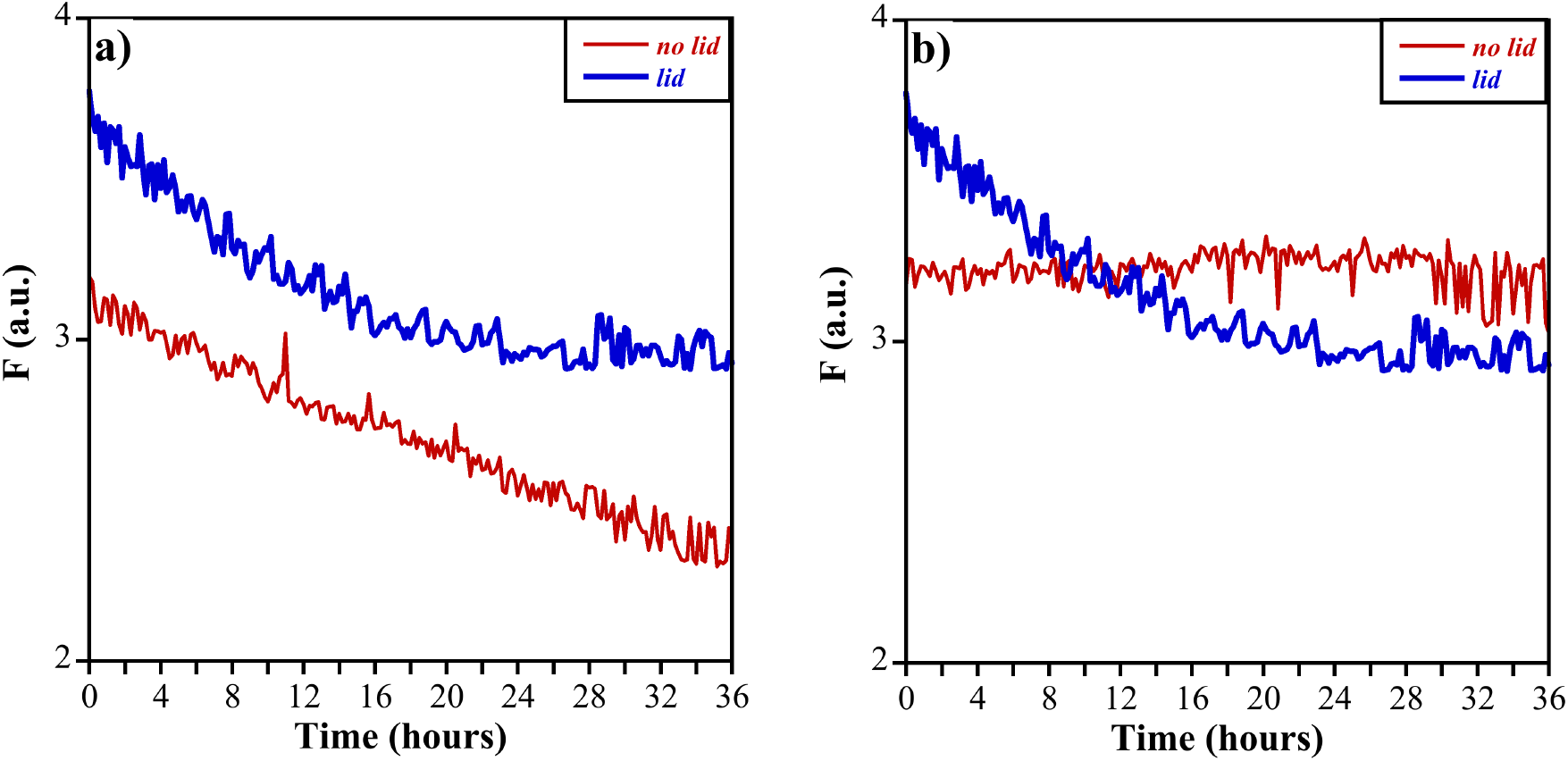
a) Focus measure dynamics for two experiments, one with a lid and one without a lid, when the microscope column is at the reference position. b) Focus measure with autofocusing and for two experiments, one with a lid and one without a lid.

The results of Figure 7 imply that autofocusing can offset the degradation in focus experienced when no autofocusing or no lid is used. However, the performance of autofocusing and that of simply using a lid are not significantly different. This would mean, *a priori*, that one could use either method. However, we submit that autofocusing might still be needed when using a lid as there are mechanisms other than evaporation that may degrade image quality. For example, as cells metabolize the media and undergo their normal life-cycle events, their refractive index can change^34^. This could cause image quality to degrade. Thus, in such a scenario, autofocusing may compensate for the loss in image quality.

### C. Autofocusing

#### Hardware

The autofocus system includes a printed circuit board (PCB) equipped with a microcontroller (Arduino Nano Every Monza, Italy) configured to control a two-phase, 400-step per rotation, stepper motor (SparkFun Electronics, Niwot, CO). The PCB has two power supply domains: an 8 V power supply for the motor and the microscope’s light source and a 5 V power supply for the microcontroller. Separating the power supplies helped protect sensitive components like the microcontroller from power supply disturbances originating from the relatively high switching currents of the stepper motor. To further enhance protection, large decoupling capacitors were used to reduce ripples on the power supplies during operation. The PCB further includes a motor driver module (DRV8833, Adafruit) that interfaces the microcontroller with the stepper motor. In addition to energizing the motor, the driver module limits the current in each phase to 1.2 A to avoid overheating.

Because fine focusing requires a low amount of torque, the stepper motor is mounted directly on the microscope. This is done using the PCB as base, without the use of brackets. Mounting slots allow for a GT-2 timing belt (2mm pitch, 6mm width from McMaster-Carr Elmhurst, IL) to provide tension adjustment. The stepper motor drives the belt using a 24T pulley (McMaster-Carr) The belt connects to the fine focus using a 92T 3D printed pulley. To withstand the high humidity of the incubator environment, the pulley was printed in NextDent Ortho clear resin (3D Systems, Rock Hill, SC). Furthermore, in order to keep the pulley from slipping around the focus knob, two 4-40 stainless screws (McMaster-Carr) were used to securly attach the pulley to the microscope’s fine focus knob. The autofocus control system is shown in Figure 8(a).

**FIG. 8.**
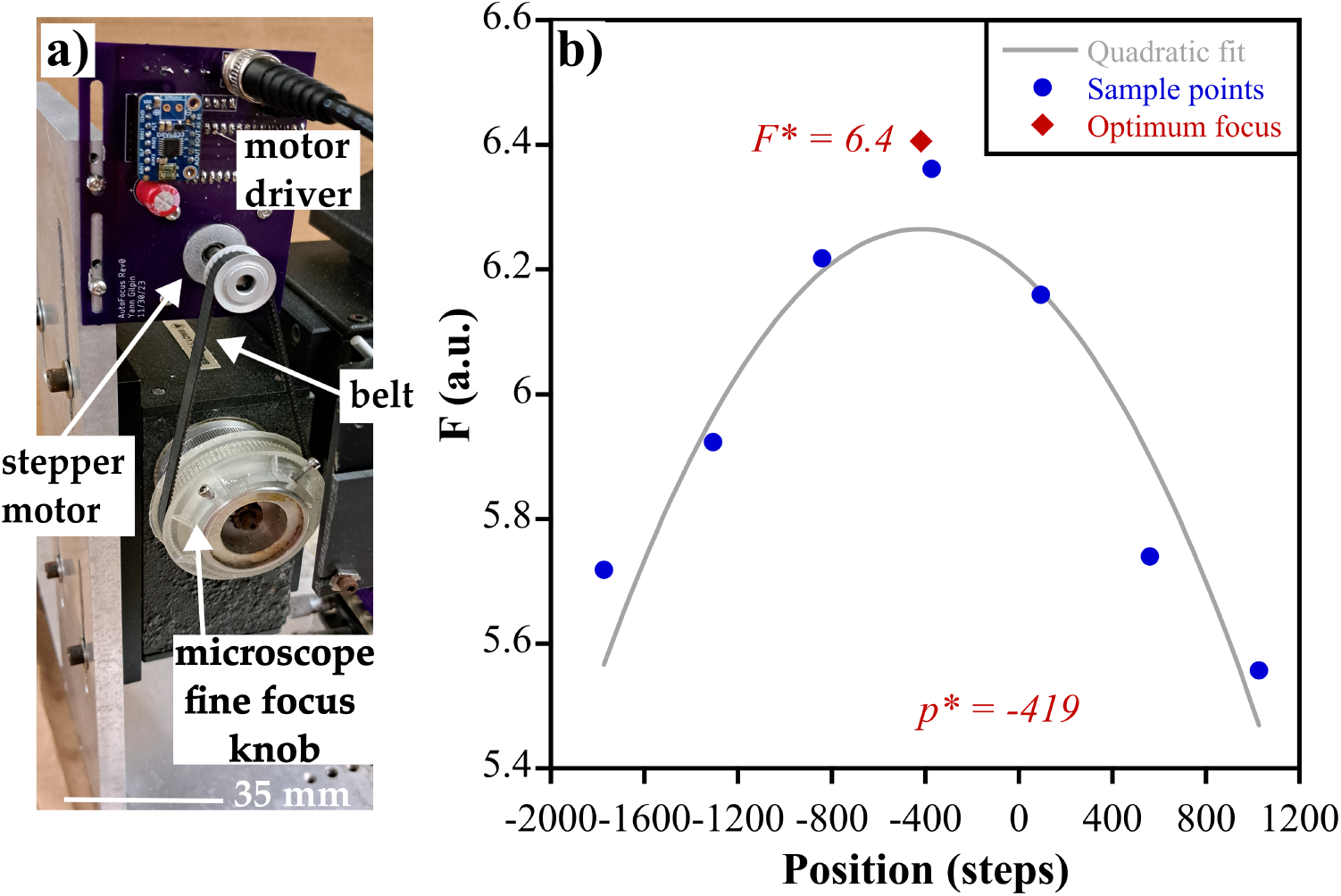
a) The autofocusing hardware include a microcontroller configured to drive a stepper motor. The motor is attached via a belt to a 3D-printed wheel affixed to the microscope’s fine focus knob. b) The autofocusing algorithm estimates the focus for 7 arbitrarily chosen positions around a current or a reference position of the microscope column. A quadratic is used to fit the focus data in the least squares sense, and the position *p*^*^ at which the quadratic is maximized. The focus at *p*^*^, *i*.*e*., the optimum focus *F*^*^, is then calculated. We note that *F*^*^ is not the maximum value of the quadratic, but rather the calculated focus measure when the microscope column is moved to the position *p*^*^.

#### Autofocusing Algorithm and Algorithm Convergence

The autofocus system’s hardware module is configured to move the microscope column and place it at an optimum position, and its software module is configured to find the optimum position based on focus measures computed from acquired images; the software module is further configured to issue commands for actuating the microscope column. The autofocus problem can be thought of as finding the position *p* for which the focus measure *F* is maximized; this is written mathematically in Equation 4.

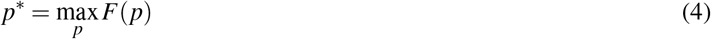

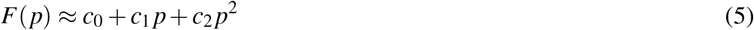

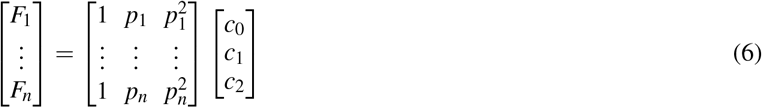

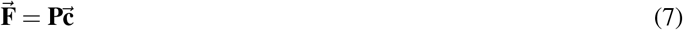

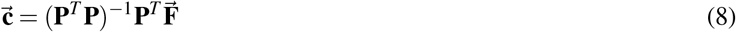

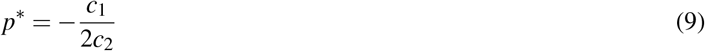

Here, *p* is the position of the microscope column, which we express in steps of the stepper motor relative to a reference position. Since *F* is a measured function rather than an analytical one, we approximate it as a quadratic polynomial for ease of optimization^35^. This is expressed in Equation 5, where *c*_0_, *c*_1_, and *c*_2_ are the real coefficients of the polynomial. To obtain the optimum position, and thus the maximum focus measure, the microscope column set sequentially to *n* distinct positions corresponding to *p*_1_, *p*_2_, …, *p*_*n*_ and focus measures *F*_1_, *F*_2_, …, *F*_*n*_ are computed for the images acquired at each of these positions. With these *n* observations, we construct the matrix equation shown in Equations 6 and 7. Because there are more observations than coefficients, Equation 7 has the least squares solution shown in Equation 8. Once the coefficients are found, the quadratic’s maximum is calculated when a goodness of fit threshold is met for the *n* observations. The position at which the maximum occurs is *p*^*^, and it can be found from the coefficients using Equation 9. Figure 8(b) shows the estimated quadratic fit for *n* = 7. The seven focus measures estimated at the *n* positions are shown, and the optimum position *p*^*^ returned by the algorithm is also noted on the graph. Once the microscope column is moved to *p*^*^, an image is acquired, and the optimum focus measure *F*^*^ is calculated.

The average convergence time *t*_*avg*_ is a key performance metric of an autofocusing system. Figure 9(a) shows a histogram of convergence times recorded for our system. The measured average convergence *t*_*avg*_ was 29.8 seconds. This result suggests that autofocusing convergence is slow for our implementation, however, because the cellular biophysical processes under study can take quite a long time (hours to days), this performance level is not detrimental to our application.

**FIG. 9.**
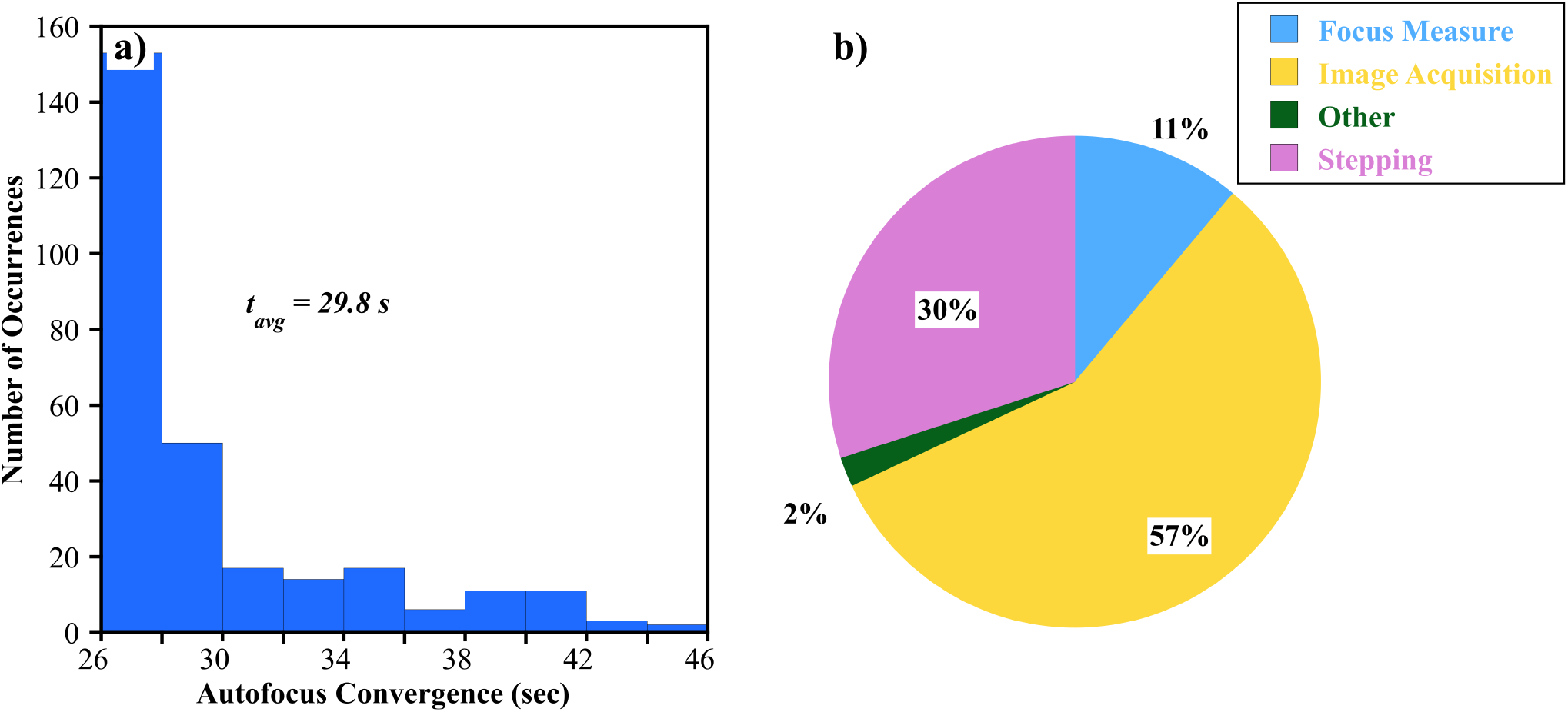
Autofocus convergence timing information for trial 1; the other trials are similar. (a) histogram of the autofocus convergence time the mean and standard deviation are 29.8s ± 4.4s (b) breakdown of the autofocus convergence with the largest slices being getting a new image and stepping to the next position.

Nevertheless, for further improvements, we show in Figure 9(b) the relative contribution to the convergence time of the different operations carried out by the software and hardware module. Image acquisition accounts for 57% of convergence time, whereas the motor’s stepping and computing the focus measures from the acquired images accounts for 30% and 11%, respectively. Additional software delays may be incurred, and they account for 2% of the convergence time.

Generally, the results of Figure 9 imply that there is a large variance in the autofocus convergence time. This variance may be caused by noise in the camera system and by vibrations in the lab, as the microscope is not placed on a vibration isolation table. Because of these factors, the autofocus control system may require additional sample points and additional images to be taken at each position. Autofocus convergence time may be traded off with image quality. This may be done using a goodness of fit measure of the quadratic as a threshold.

### D. Cell Counting

Cell counting with a computer vision algorithm helps correlate measured capacitance time series data with cell culture growth. For instance, Smith *et al*. showed that the average capacitance measured across 16 capacitance-sensing electrodes had a strong correlation with the evolution of cell numbers in a region of interest^11^. From this observation, they developed a temporal model that related average measured capacitance with cell numbers. However, the computer vision algorithm’s performance strongly depended on image quality and, as a corollary, on the ability to maintain a high focus measure. The results featured in this section show that using a lid and/or autofocusing improves the performance of computer vision-based cell detection and cell counting software. Our study used a computer vision algorithm previously developed in our group, which we briefly describe below, but a more in-depth coverage can be found in refs.^11,12^.

Compared to state-of-the-art methods, which train with focused images including background features and references such as electrode or interconnect patterns^36,37^, our computer vision algorithm averages these artifacts out by using a correlation filter across multiple labeled images^38^. Generally, our algorithm first includes processing the images after acquisition to enhance quality. This step can include sharpening and/or applying a false color scheme to enhance the contrast between the cells and the background. Next, a reference image is used to build a template that matches cellular features. During execution, the algorithm matches the template with image regions *F*_*i*_, returning a correlation measure between the template and images features in a specific region^39,40^. Locations with high correlations are flagged as likely having cells present^41^; in Figures 10 and 11, markers for high correlation are shown as magenta-colored dots. To be effective, the template is constructed as a correlation filter that must be trained with labeled features that are known *a priori* to correspond to cell features. The filter training process optimizes a convolution response *G*_*i*_ between input patches *F*_*i*_ in a training set and the to-be-trained filter *H*. This training process is formulated in Equation 10.

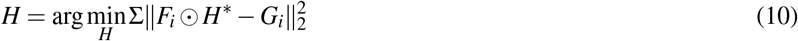

**FIG. 10.**
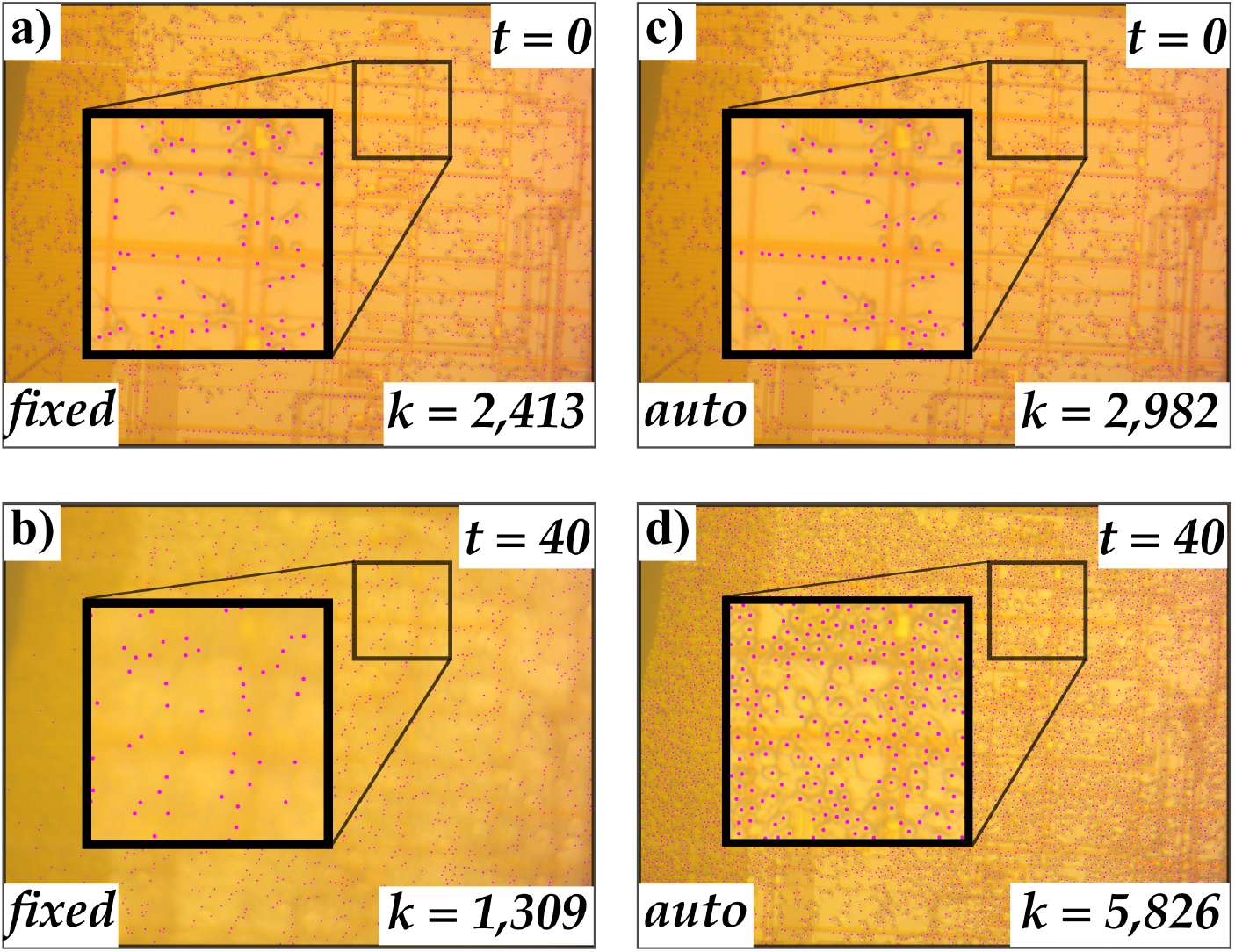
Cell counting with no lid. (a,b) The microscope column is held at the reference position, *i*.*e*., at the fixed focus position, which is set at the beginning of the experiment (*t* = 0 hours). The number of cells counted (including false positives) was *k* = 2, 413 in the image. At time *t* = 40 hours, the image degrades due to media evaporation. Consequently, the cell counting code can only identify *k* = 1, 309 objects in the field of view. (c, d) At *t* = 0 hours, the microscope is held the reference position, and we registered *k* = 2, 982 objects. With autofocusing, at time *t* = 40 hours, high image quality is maintained, and *k* = 5, 826 objects are counted. We note that here, the cells are in a region of interest that is sparsely populated by interdigitated capacitance-sensing electrodes.^11^.

**FIG. 11.**
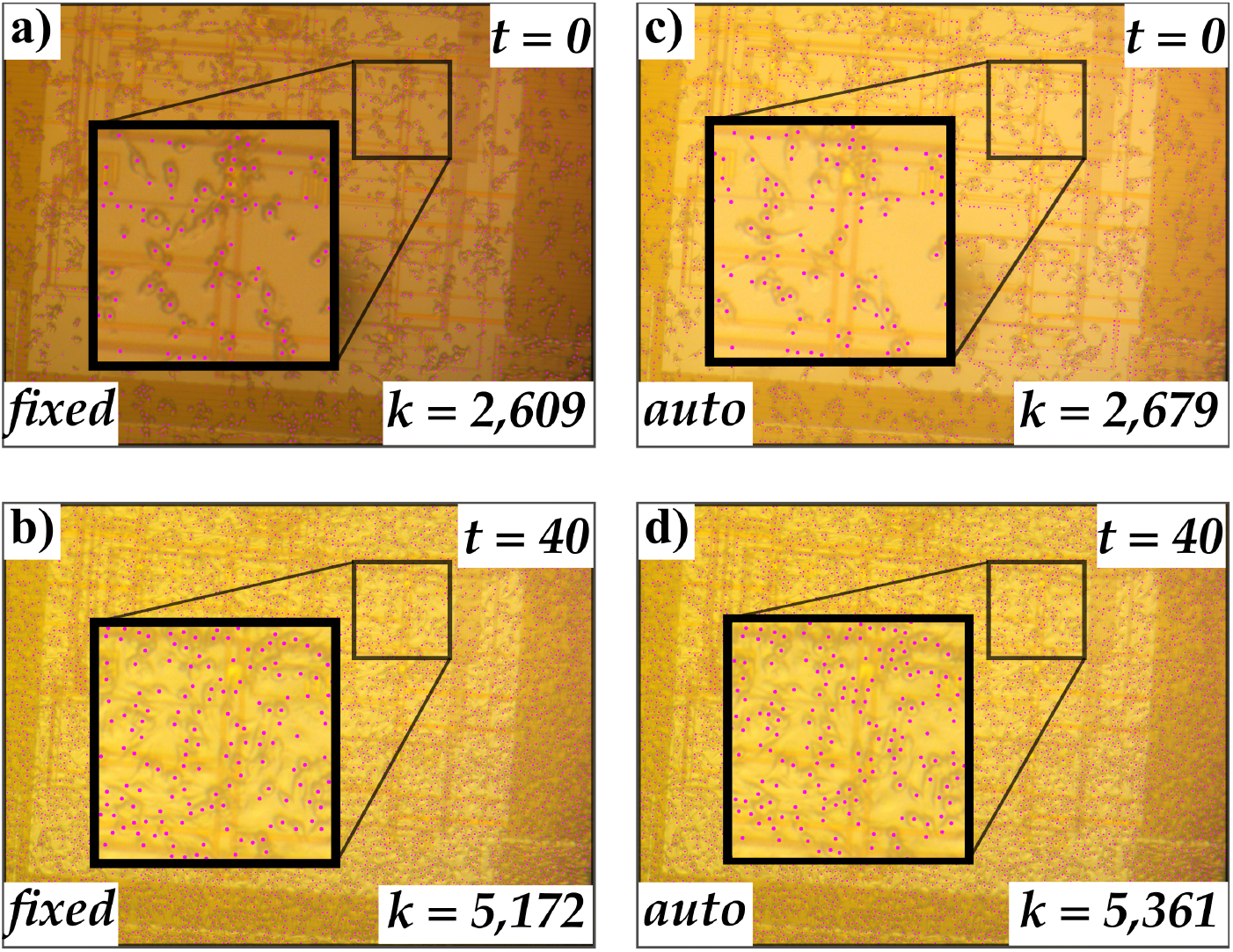
Cell counting with an immersion lid. (a,b) The microscope column is held at the reference position, *i*.*e*., at the fixed focus position, which is set at the beginning of the experiment (*t* = 0 hours). The number of cells counted (including false positives) was *k* = 2, 609 in the image. At time *t* = 40 hours, image quality is maintained. The computer vision code identifies *k* = 5, 172 objects in the field of view. (c, d). With autofocusing, at time *t* = 0 hours, *k* = 2, 679 and at *t* = 40 hours, *k* = 5, 361. With autofocusing, image quality is maintained throughout the experiment. and the counts are consistent with the fixed focus experiment. Again, we note that here, the cells are in a region of interest that is sparsely populated by interdigitated capacitance-sensing electrodes.^11^.

The algorithm was evaluated as a standard object detection task using a manually labeled data set, and the average precision was found to be 0.871 for an intersection over union (IoU) set at 50%^42^. We note that precision, *i*.*e*., the ability to correctly detect a cell, also depends on the intrinsic features of the images under test and the experimental conditions during image acquisition. In the former case, the chip’s top metal lines, though helpful when calculating a focus measure, can lead to false positives in cell counting. In the latter case, vibrations, errors in positioning, and random variations in the light field can also lead to false positives^43^, which introduces some variance in the cell counts estimated. Nevertheless, our scope in this work is concerned with studying the interplay between, evaporation dynamics, autofocusing, and cell detection. As such, we report a total number of estimated objects in the field of view, denoted *k*, and this number accounts for cells successfully flagged by the algorithm, as well as false positives.

As a measure of autofocusing performance, we compare the value of *k* at the beginning and at the end of each experiment, under various focus conditions (autofocusing and fixed focusing), and under two evaporation conditions (lid and no lid). Figures 10(a) and (b) respectively show the beginning and endpoint for an experiment with no lid and no autofocusing. At time t = 0, the focus is fixed manually and the microscope column is held at the reference position. At time t = 40, the focus degrades as a result of evaporation. This can be seen from the blur in Figure 10(b). For the two cases, the computer vision algorithm returns *k* = 2, 413 and *k* = 1, 309 respectively. With the blurred image, the computer vision algorithm is unable to register all the cells present in the field of view. In contrast, with autofocusing, the cells can be clearly resolved by the computer vision algorithm at the endpoint. For example, Figure 10(c) reports *k* = 2, 982 at the reference position and *k* = 5, 826 at the endpoint. This means that autofocusing maintains imaging quality across the whole experiments, allowing proper estimation of the number of cells present in the field of view.

Figures 11(a) and (b) respectively show the beginning and endpoint of another experiment in which a lid was used but without autofocusing. Both images are clear, which means that a high focus was maintained by the autofocusing algorithm. At the start of the expriment, the computer-vision code registered 2,609 objects and 5,172 objects at the endpoint. Figures 11(c) and (d) show similar numbers and the same high-quality images at both start and finishing when autofocusing was used actively. This means that autofocusing achieves the same results as when a lid is used without autofocusing.

## IV. DISCUSSION AND CONCLUSIONS

This paper presented a quantitative study of image quality degradation as a function of cell media evaporation in an in-incubator lab-on-CMOS capacitance sensing platform. We showed that evaporation still occurs when a novel immersion lid is used, although at a smaller rate than when no lid is used. The immersion lid was provisioned with side ports to allow gas exchange with the cell media, allowing the cells to thrive. We showed that this lid allows cell culture for up to 40 hours with high-quality images. In addition, the immersion lid has the advantage of reducing the evaporation rate. This is an important feature for cell culture, as a reduced evaporation rate means that the cell media’s respective concentrations of salts, proteins, and small molecules do not experience large changes over time, which allows for uniform culture conditions over long periods.

We showed that autofocusing was another viable mechanism for ensuring high-quality image acquisition. To do so, we constructed a custom autofocusing system (hardware and software) with which we retrofitted a standard upright optical microscope. Combining the immersion lid with autofocusing did not yield better results than when one technique was used without the other. We also note that both methods have shortcomings. The main drawback of the immersion lid is maintaining cleanliness, as cleaning involves several process steps. Moreover, the auto-focusing system increases the amount of time that the microscope light is on; this may be detrimental to cell cultures. This is especially important if the microscope uses a halogen or an incandescent light source; in our experience, these types of sources can generate enough heat to cause local temperature gradients in the cell culture and compromise cell viability. Furthermore, the slow convergence speed of the auto-focusing system may be improved using a faster image acquisition system, as we found that image acquisition was the main impediment to a fast auto-focusing convergence time.

Lastly, we note that a better microscope, higher magnification, or an immersion lens could undoubtedly mitigate some of the issues raised in this paper in regards to focus degradation and imaging quality in general. Moreover, a perfusion system could also be used to mitigate cell media evaporation. However, we caution that, by design, the imaging hardware overhead in the present study was chosen to be minimal to ensure simplicity. And, also for the sake of simplicity, we did not consider the use of a perfusion system. These design choices were made because, at scale, we do not envision calibrating our lab-on-CMOS sensors with complex microscopy hardware and perfusion systems, but rather with simple upright bright-field microscopy hardware having long working distances and low magnification so that a wide area of a chip in a culture well may be imaged. Above all, our vision is that the imaging system itself can also be miniaturized and integrated on a scanning rack inside the incubator to calibrate a large number of sensors deployed in a well plate format^8^ simply by positioning an imaging head over a specific sensor. In sum, despite the above-noted limitations, our results represent the first quantitative study to highlight the challenges encountered when calibrating lab-on-CMOS microsystems using microscopy as ground-truth data. As such, they lay the ground work for more robust calibration techniques to be developed.

## ACKNOWLEDGMENTS

This article is based on work supported by the National Science Foundation under Grant No. 2442346, and any opinions, findings, conclusions, or recommendations expressed in this article are those of the authors and do not necessarily reflect the views of the National Science Foundation. We further thank Pamela Abshire, Bathiya Senevirathna, and Sheung Lu for sensor design. Lastly, we thank Zixin Chen for help with cell culture.

## AUTHOR DECLARATIONS

### Conflict of Interest

The authors have no conflicts to disclose.

### Author Contributions

Y.G. designed the experiments, the auto-focusing system, the immersion lid, and he developed the integrative packaging method, and he designed the daughter board. C.-Y. L. developed analytical methods for image processing, and he designed the information infrastructure for cataloging measured data. Further, C.-Y. L. designed the mother board. He also developed the code for data acquisition and control. Y.G. and M.D. designed the study and its conceptual aspects; M.D. supervised Y.G. and C.-Y. L., secured funding, and provided resources for executing the study. All authors contributed to writing the article and approved the submitted version.

## DATA AVAILABILITY

The data that support the findings of this study are available from the corresponding author upon reasonable request.

## Notes

### Competing Interest Statement

The authors have declared no competing interest.

## REFERENCES

1 M. Jeyaraman, N. Jeyaraman, S. Ramasubramanian, K. P. Iyengar, and V. K. Jain, “Lab-on-chip technology for clinical and medical application,” in Advanced Sensors for Smart Healthcare (Elsevier, 2025) pp. 287–297.

2 Y. H. Ghallab and Y. Ismail, “CMOS based lab-on-a-chip: Applications, challenges and future trends,” IEEE Circuits and Systems Magazine 14, 27–47 (2014).

3 C. D. Chin, V. Linder, and S. K. Sia, “Commercialization of microfluidic point-of-care diagnostic devices,” Lab on a Chip 12, 2118 (2012).

4 M. Mauk, J. Song, H. H. Bau, R. Gross, F. D. Bushman, R. G. Collman, and C. Liu, “Miniaturized devices for point of care molecular detection of HIV,” Lab on a Chip 17, 382–394 (2017).

5 N. Lu, H. M. Tay, C. Petchakup, L. He, L. Gong, K. K. Maw, S. Y. Leong, W. W. Lok, H. B. Ong, R. Guo, K. H. H. Li, and H. W. Hou, “Label-free microfluidic cell sorting and detection for rapid blood analysis,” Lab on a Chip 23, 1226–1257 (2023).

6 M. Toner and D. Irimia, “Blood-on-a-Chip,” Annual Review of Biomedical Engineering 7, 77–103 (2005).

7 W. Jung, J. Han, J.-W. Choi, and C. H. Ahn, “Point-of-care testing (POCT) diagnostic systems using microfluidic lab-on-a-chip technologies,” Microelectronic Engineering 132, 46–57 (2015).

8 S. Chitale, W. Wu, A. Mukherjee, H. Lannon, P. Suresh, I. Nag, C. M. Ambrosi, R. S. Gertner, H. Melo, B. Powers, H. Wilkins, H. Hinton, M. Cheah, Z. G. Boynton, A. Alexeyev, D. Sword, M. Basan, H. Park, D. Ham, and J. Abbott, “A semiconductor 96-microplate platform for electrical-imaging based high-throughput phenotypic screening,” Nature Communications 14, 7576 (2023).

9 K. Hu, C. E. Arcadia, and J. K. Rosenstein, “A Large-Scale Multimodal CMOS Biosensor Array With 131,072 Pixels and Code-Division Multiplexed Readout,” IEEE Solid-State Circuits Letters 4, 48–51 (2021).

10 R. A. Croce, S. Vaddiraju, A. Legassey, K. Zhu, S. K. Islam, F. Papadimitrakopoulos, and F. C. Jain, “A Highly Miniaturized Low-Power CMOS-Based pH Monitoring Platform,” IEEE Sensors Journal 15, 895–901 (2015).

11 K. Smith, C.-Y. Lin, Y. Gilpin, E. Wayne, and M. Dandin, “Measuring and modeling macrophage proliferation in a lab-on-CMOS capacitance sensing microsystem,” Frontiers in Bioengineering and Biotechnology 11 (2023), 10.3389/fbioe.2023.1159004.

12 C.-Y. Lin and M. Dandin, “Machine Learning Identification and Classification of Mitosis and Migration of Cancer Cells in a Lab-on-CMOS Capacitance Sensing Platform,” IEEE Journal of Biomedical and Health Informatics 29, 1504–1513 (2025).

13 B. Senevirathna, A. Castro, M. Dandin, E. Smela, and P. Abshire, “Lab-on-CMOS capacitance sensor array for real-time cell viability measurements with I2C readout,” in 2016 IEEE International Symposium on Circuits and Systems (ISCAS) (IEEE, 2016) pp. 2863–2866.

14 S. B. Prakash and P. Abshire, “On-Chip Capacitance Sensing for Cell Monitoring Applications,” IEEE Sensors Journal 7, 440–447 (2007).

15 C. L. Davey and D. B. Kell, “The low-frequency dielectric properties of biological cells,” in Bioelectrochemistry of Cells and Tissues (Birkhäuser Basel, Basel, 1995) pp. 159–207.

16 M. S. R. Sajal, K.-C. Lin, Y. Gilpin, V. Walawalkar, F. Dehghandehnavi, R. E. Taylor, and M. Dandin, “Towards CMOS Capacitance Sensors for DNA Origami Characterization,” in 2023 International Electron Devices Meeting (IEDM) (IEEE, 2023) pp. 1–4.

17 S. Forouhi, R. Dehghani, and E. Ghafar-Zadeh, “CMOS based capacitive sensors for life science applications: A review,” Sensors and Actuators A: Physical 297, 111531 (2019).

18 B. P. Senevirathna, S. Lu, M. Dandin, J. Basile, E. Smela, and P. A. Abshire, “Real-Time Measurements of Cell Proliferation Using a Lab-on-CMOS Capacitance Sensor Array,” IEEE Transactions on Biomedical Circuits and Systems 12, 510–520 (2018).

19 S. Lu, B. Senevirathna, M. Dandin, E. Smela, and P. Abshire, “System Integration of IC chips for Lab-on-CMOS Applications,” in 2018 IEEE International Symposium on Circuits and Systems (ISCAS) (IEEE, 2018) pp. 1–5.

20 B. P. Senevirathna, S. Lu, M. P. Dandin, E. Smela, and P. A. Abshire, “Correlation of Capacitance and Microscopy Measurements Using Image Processing for a Lab-on-CMOS Microsystem,” IEEE Transactions on Biomedical Circuits and Systems 13, 1214–1225 (2019).

21 B. Senevirathna, S. Lu, M. Dandin, J. Basile, E. Smela, and P. Abshire, “High resolution monitoring of chemotherapeutic agent potency in cancer cells using a CMOS capacitance biosensor,” Biosensors and Bioelectronics 142, 111501 (2019).

22 B. Senevirathna, S. Lu, E. Smela, and P. Abshire, “An Imaging Platform for Real-Time In Vitro Microscopic Imaging for Lab-on-CMOS Systems,” in 2019 IEEE Biomedical Circuits and Systems Conference (BioCAS) (IEEE, 2019) pp. 1–4.

23 N. Halonen, J. Kilpijärvi, M. Sobocinski, T. Datta-Chaudhuri, A. Hassinen, S. B. Prakash, P. Möller, P. Abshire, S. Kellokumpu, and A. Lloyd Spetz, “Low temperature co-fired ceramic packaging of CMOS capacitive sensor chip towards cell viability monitoring,” Beilstein Journal of Nanotechnology 7, 1871–1877 (2016).

24 J. Kilpijärvi, N. Halonen, M. Sobocinski, A. Hassinen, B. Senevirathna, K. Uvdal, P. Abshire, E. Smela, S. Kellokumpu, J. Juuti, and A. Lloyd Spetz, “LTCC Packaged Ring Oscillator Based Sensor for Evaluation of Cell Proliferation,” Sensors 18, 3346 (2018).

25 M. Dandin, I. D. Jung, M. Piyasena, J. Gallagher, N. Nelson, M. Urdaneta, C. Artis, P. Abshire, and E. Smela, “Post-CMOS packaging methods for integrated biosensors,” in Proceedings of IEEE Sensors (2009).

26 Y. Huang and A. J. Mason, “Lab-on-CMOS integration of microfluidics and electrochemical sensors,” Lab on a Chip 13, 3929 (2013).

27 Y. Gilpin, C.-Y. Lin, M. Forssell, S. Zheng, P. Grover, and M. Dandin, “Tracking the Effects of Tumor Treating Fields on Human Breast Cancer Cells in vitro Using a Capacitance Sensing Lab-on-CMOS Microsystem,” in 2022 29th IEEE International Conference on Electronics, Circuits and Systems (ICECS) (IEEE, 2022) pp. 1–4.

28 J. Pech-Pacheco, G. Cristobal, J. Chamorro-Martinez, and J. Fernandez-Valdivia, “Diatom autofocusing in brightfield microscopy: a comparative study,” Proceedings 15th International Conference on Pattern Recognition 3, 314–317 (2000).

29 W. Boecker, W. Rolf, W.-U. Muller, and C. Streffer, “Autofocus algorithms for fluorescence microscopy,” in Proc. SPIE 2847, Applications of Digital Image Processing XIX, edited by A. G. Tescher (1996) pp. 445–456.

30 F. C. A. Groen, I. T. Young, and G. Ligthart, “A comparison of different focus functions for use in autofocus algorithms,” Cytometry 6, 81–91 (1985).

31 E. Krotkov, “Focusing,” International Journal of Computer Vision 1, 223–237 (1988).

32 M. Subbarao, “Focusing techniques,” Optical Engineering 32, 2824 (1993).

33 T. Yeo, S. Ong Jayasooriah, and R. Sinniah, “Autofocusing for tissue microscopy,” Image and Vision Computing 11, 629–639 (1993).

34 M. A. Calin, M. R. Calin, and C. Munteanu, “Determination of the complex refractive index of cell cultures by reflectance spectrometry,” The European Physical Journal Plus 129, 116 (2014).

35 J.-M. Geusebroek, F. Cornelissen, A. W. Smeulders, and H. Geerts, “Robust autofocusing in microscopy,” Cytometry 39, 1–9 (2000).

36 Z. Zou, K. Chen, Z. Shi, Y. Guo, and J. Ye, “Object Detection in 20 Years: A Survey,” Proceedings of the IEEE 111, 257–276 (2023).

37 O. Ronneberger, P. Fischer, and T. Brox, “U-Net: Convolutional Networks for Biomedical Image Segmentation,” (2015) pp. 234–241.

38 C.-Y. Lin, Y. Gilpin, Z. Chen, E. Wayne, and M. Dandin, “Towards a Lab-on-a-Chip as a Service (LoCaaS) Framework for in Vitro Cancer Cell Assays,” in 2024 22nd IEEE Interregional NEWCAS Conference (NEWCAS) (IEEE, 2024) pp. 268–272.

39 D. Bolme, J. R. Beveridge, B. A. Draper, and Y. M. Lui, “Visual object tracking using adaptive correlation filters,” in 2010 IEEE Computer Society Conference on Computer Vision and Pattern Recognition (IEEE, 2010) pp. 2544–2550.

40 J. F. Henriques, R. Caseiro, P. Martins, and J. Batista, “High-Speed Tracking with Kernelized Correlation Filters,” IEEE Transactions on Pattern Analysis and Machine Intelligence 37, 583–596 (2015).

41 A. Neubeck and L. Van Gool, “Efficient Non-Maximum Suppression,” in 18th International Conference on Pattern Recognition (ICPR’06) (IEEE, 2006) pp. 850–855.

42 R. Padilla, S. L. Netto, and E. A. B. da Silva, “A Survey on Performance Metrics for Object-Detection Algorithms,” in 2020 International Conference on Systems, Signals and Image Processing (IWSSIP) (IEEE, 2020) pp. 237–242.

43 L. Yuan, T. Guo, Z. Qiu, X. Fu, and X. Hu, “An Analysis of the Focus Variation Microscope and Its Application in the Measurement of Tool Parameter,” International Journal of Precision Engineering and Manufacturing 21, 2249–2261 (2020).

